# Global patterns of taxonomic and functional diversity in hot springs microbiomes

**DOI:** 10.1101/2025.08.04.668570

**Authors:** José Ignacio Arroyo, Beatriz Díez, Pablo A. Marquet

## Abstract

A reformulation of the metabolic theory of ecology has been proposed based on a derivation of the Eyring-Polanyi equation, leading to a reduced model with the form of a power-law with exponential cut-off (PLECO). Moreover, the recent integration of this new model and metagenomic DNA sequencing theory has led to the hypothesis that, given that temperature affects cell size and assemblage abundance, it should also affect the number of genes, metabolisms, and populations in a microbial community. In particular, for microbial communities, the model predicts that the diversity of taxa, their genes, and metabolisms should respond to temperature according to the PLECO model, and after scaling the data using the estimated parameters, all data should collapse into a single curve. To test these predictions, we use global hot springs microbial communities; geothermally heated environments that include aquatic and terrestrial habitats with large temperature gradients. We analyzed a global database (five continents) of 19 hot springs metagenomes covering a wide gradient of temperature (32-90°C). We found that the alpha diversity of taxa, genes, and their metabolisms decreases with temperature, in agreement with the predictions of the PLECO model. Further, after non-dimensionalization of the model, we derive a single nondimensional curve for taxa and functions. Finally, using general linear models, we analyzed the effects of pH, latitude, and substrate (soil, water, or both), in addition to temperature, to explain the observed diversity pattern. We find that they explain a lower proportion of the variance, but pH can explain up to half of the variance in global diversity in these ecosystems, indicating the need to include pH in the theory. Our study supports the hypothesis that temperature regulates the structure of community genomes.

## Introduction

The metabolic theory of ecology (MTE) integrates the universal constraints imposed by size and temperature on biological rates across levels of biological organization, from organisms to ecosystems (Brown et al. 2004, West et al. 2005, Sibly et al. 2012). This framework emerged from the integration of Kleiber’s law, relating body size and metabolic rate, and the empirical Arrhenius equation, relating temperature and the rate of (bio)chemical reactions (Brown et al. 2004). This model was extended to predict different ecological processes from individuals to ecosystems (e.g., Allen et al., Brown et al. 2004). Despite the broad scope of predictions of the MTE, there are still challenges, particularly regarding the integration with other theories and the prediction of new patterns and driving variables. For instance, at the community level, theoretical predictions for species richness (Allen et al. 2002) and abundance of heterotrophs (Allen et al. 2007) have been derived assuming that other aspects of ecological complexity (e.g., phylogenetic and functional; Naeem et al. 2016) are not important, which is at odds with theories and data showing that they may be critical to predicting ecological stability and ecosystem functioning (Thébault and Loreau 2003, Thébault and Fontaine 2011).

We have recently suggested the use of the Eyring-Evans-Polanyi equation, instead of Arrhenius’equation, in predicting biodiversity patterns on the grounds that the former is grounded on solid thermodynamic arguments and is widely accepted as a descriptor of the role of temperature on chemical reactions (Arroyo et al. 2022). This equation is

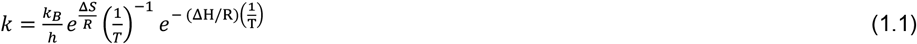

where *k* is reaction rate (s^-1^, for a first-order reaction), *k*_*B*_ is the Boltzmann constant, *h* is the Plank constant (= 6.62 ×10-34 Js^-1^), *T* is temperature (K), ΔS is the entropy change (J/K), ΔH enthalpy change (J), *R* is the gas constant (= 8.31 JK^-1^ mol^-1^). Using this equation, we have proposed a general model of temperature dependence for biological rates, which yields an equation with the form of a power law with an exponential cut-off

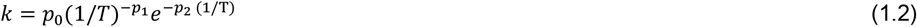

Here, *k* could be the number of species (taxa), genes of groups of related genes (gene families, paths or modules), 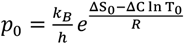, *p*_1_ = (ΔG/R)+1, and *p*_2_ = ΔH/R, *T* is temperature is in Kelvin, *R* is the gas constant, ΔH is free enthalpy change (J), ΔS_0_ entropy at T_0_, ΔC is the heat capacity change. Negative values of ΔC gives an asymmetric concave curve that accounts for the breakpoint commonly observed in all temperature response curves. A second prediction of our models is that scaling the data using estimated parameters should give a single curve according to

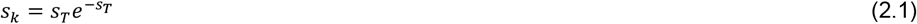

Where 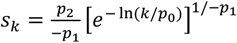 and 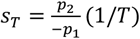 Alternatively, scaling can be done as

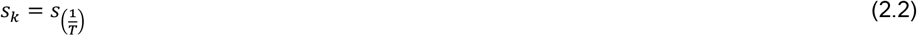

Where 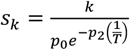 and 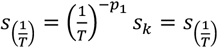 Eq. 2.1 predicts that data should collapse into a curved pattern, whereas Eq. 2.2 predicts that data should collapse into a linear pattern.

After suggesting this new model of temperature dependence, we integrated metabolic ecology and DNA shotgun sequencing theory based on the empirical relationships between temperature and cell size and the effect of mean cell size on assemblage abundance, thus implying that temperature affects assemblage abundance. Then we simplified DNA shotgun sequencing predictions for taxa and gene reads by integrating the genome size-cell size, and population abundance and cell size relationships to predict that taxa and gene reads depend uniquely on population and assemblage abundance, respectively. Finally, by assuming that the abundance and number of genes and groups of genes scale with assemblage abundance, we predicted that ultimately, temperature affects the number of genes and groups of related genes of a community. This results in specific testable predictions of temperature dependence in the richness of taxa, their genes, and metabolisms according to Eq. 1.2 The predictions were relatively well supported by genomes and metagenomes from different taxa and ecosystems, including data from human, terrestrial, and aquatic microbiomes (Chapter 2).

A natural laboratory where these predictions can be further tested is hot springs, where are founds temperature gradients are found along a few meters (see for example Alcamán et al. 2015). These ecosystems are geothermally heated patches, with temperatures higher than the mean environmental temperature (Pentecost et al. 2003, Pentecost et al. 2005, O’Gorman et al. 2014), and where it is often possible to find microbial mats in their surroundings. Comparatively, hot spring microbial mats are less diverse than other terrestrial (e.g., soil) and marine and freshwater ecosystems (Lozupone and Knight 2007, Aanniz et al. 2015) but contain representatives of most Archaeal and Bacterial lineages (Sharp et al. 2014). These communities can be found both as components of the pelagic environment in the water column or forming mats attached to the bottom sediments and other substrates; the latter are often more diverse than pelagic ones (Wang et al. 2014). Most studies examining ecological patterns in hot springs have been conducted at larger scales. As a general pattern, studies at the scales of meters have shown that abundance and diversity and composition patterns are explained by the temperature, pH and geochemistry with geography becoming important at kilometer scales (Miller et al. 2009, Ionescu et al. 2010, Cuecas et al. 2011, Tobler et al. 2011, Everroad et al. 2012, Cole et al. 2013, Wang et al. 2013, Inskeep et al 2013, Headd et al. 2014, Herbold et al. 2014, Valverde et al. 2014, Sharp et al. 2014, Wang et al. 2014; Menzel et al 2015; Power et al 2018, Arroyo et al. 2025).

Here, we fitted four predictions of our recently developed framework of metabolic ecology and metagenomics on global hot springs metagenomes; in particular we predict a temperature dependence of i) the number of taxa, ii) the number of genes, and iii) the number of groups of related genes (e.g. metabolic pathway) and iv) scaling the data with the estimated parameters should result in a single curve according to eqs. 2.1 and 2.2. v). In addition, we also assessed the effects of pH, latitude, and type of substrate as predictors of taxonomic and functional diversity as they have been shown to explain diversity in these and other ecosystems (Fierer and Jackson 2006, Fuhrman et al. 2008, Inskeep et al. 2013).

## Results

We begin by fitting the relationship between the number of species and temperature. The curve model (eq. 1.2) had a significant fit (Figure 1a), although its low curvature was statistically different from the linear model of eq. 1.1 (ΔAIC=1.995). Then we fitted the relationship between the number of genes and groups of related genes (e.g., families) with temperature. For this, we used reads annotated according to the functional classification of the SEED database with three different levels of resolution: SEED level 1 (corresponding to biological process), SEED level 2 (corresponding to metabolic pathways, gene families), and SEED level 3 (corresponding to genes, see Methods). The number of genes (SEED level 3) (Fig. 1) and groups of related genes (SEED level 2 and SEED level 1) (Fig. 2c and Fig. 2d, respectively) decreased significantly with temperature. The number of genes and ‘subsystems’ all fit well to the PLECO model of Eq. 1.2 with an explained variance that varied between 41 to 45%. (Table 1). The estimated parameters of the model (Eq. 1.2) consistently decreased with the level of organization from genes to taxa (Table 1). We also fitted equations. 2.1 and 2.2 to test the universality of temperature dependence regardless of the ecological variable (taxonomic or functional richness). As predicted, data collapsed into a single dimensionless curve according to Eq. 2.1 (Figure 2) or line according to Eq. 2.2 (Figure 2, left lower corner).

**Fig. 1.**
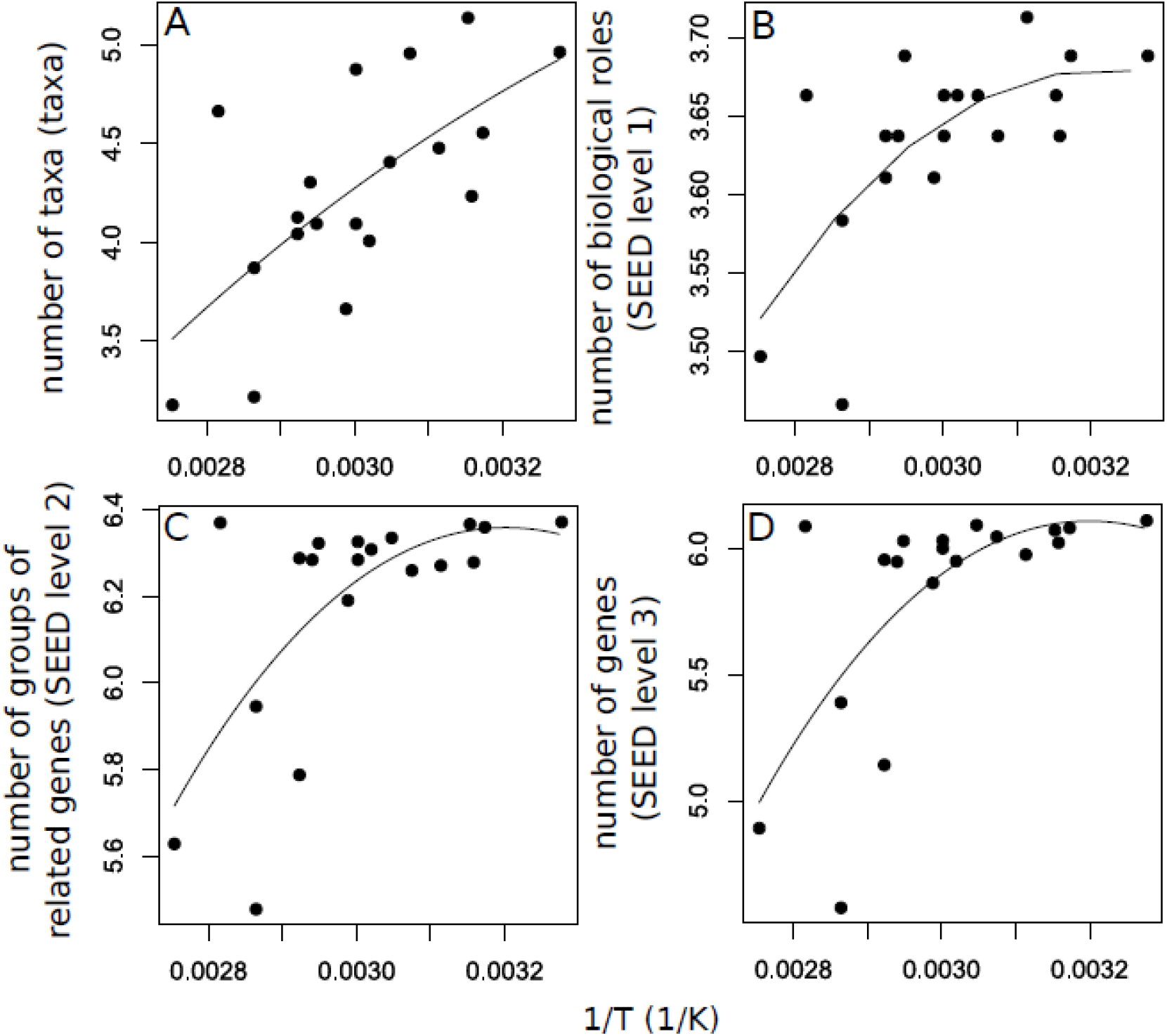
Relationship between (a) ln (taxa richness), (b) ln (SEED level 1; biological roles), (c) ln (SEED level 2; families, pathways) and (d) ln(SEED level 3; genes) and 1/T. In the four cases there was a decrease with temperature (or increases with the inverse of temperature, 1/T) in the temperature range 32-90°C.

**Fig. 2.**
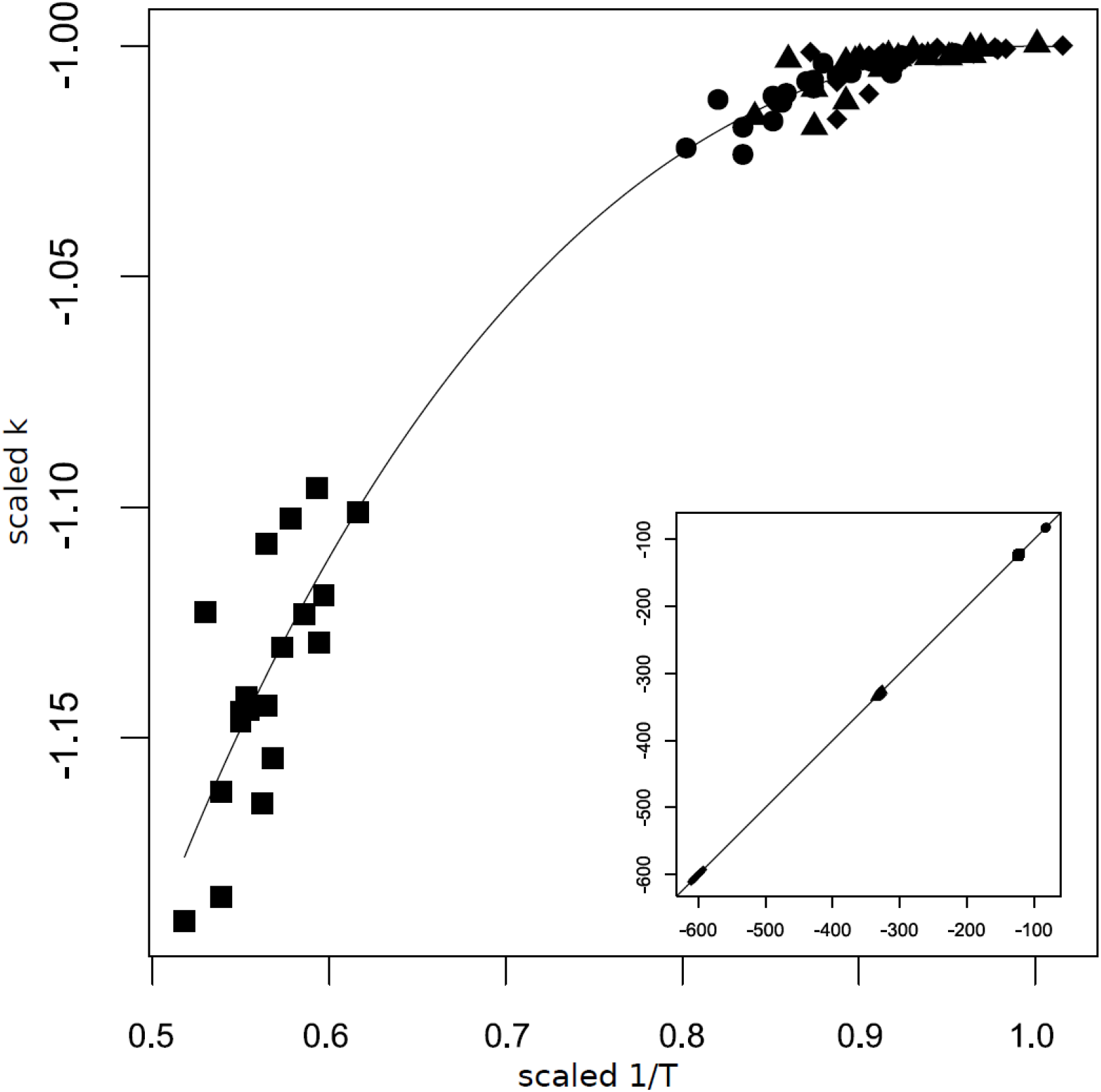
Universal temperature dependent curve for taxa and functions. Nondimensionalization based on scales rate and temperature (see Results) showing how temperature-dependence data regardless of variable data collapses into a single curve. Main panel corresponds to Eq. 2.1 and lower left panel to Eq. 2.2. (◼) Taxa richness, (•) SEED level 1 (biological roles), (▲) SEED level 2 (metabolic pathways, gene families), (⧫) SEED level 3 (genes).

**Table 1.** Estimated parameters of models implemented for the effects of temperature (T) (Fig. 1), pH and habitat type (with two levels: terrestrial and aquatic; H).

| Eq. 1.2 | p<sub>0</sub> | p<sub>1</sub> | p<sub>2</sub> | r-squared | p-value |
| --- | --- | --- | --- | --- | --- |
| Taxa-T | 91.23 | 14.04 | -1809.72 | 0.45 | 0.008 |
| SEED 1-T | 113.26 | 16.22 | -5126.67 | 0.41 | 0.01 |
| SEED 2-T | 427.3 | 62.39 | -19543.26 | 0.44 | 0.009 |
| SEED 3-T | 652.17 | 95.83 | -29858.97 | 0.43 | 0.01 |
| Linear regressions | Intercept | Slope |  |  |  |
| Taxa-pH | 3.47 | 0.11 |  | 0.14 | 0.11 |
| SEED 1-pH | 3.46 | 0.02 |  | 0.49 | 0.0007 |
| SEED 2-pH | 5.42 | 0.11 |  | 0.57 | 0.0001 |
| SEED 3-pH | 4.52 | 0.19 |  | 0.57 | 0.0001 |
| Taxa-latitude | 4.11 | 0.009 |  | 0.1 | 0.24 |
| SEED 1-latitude | 3.64 | 0.0003 |  | 0.04 | 0.46 |
| SEED 2-latitude | 6.23 | 0.02 |  | 0.29 | 0.03 |
| SEED 3-latitude | 5.97 | 0.001 |  | 0.13 | 0.16 |

In addition to temperature, we studied the effects on taxonomic and functional diversity of other environmental variables that were recorded in compiled data and our sampling: pH and latitude. Both the logarithm of the number of taxa, genes, or groups of genes and pH fitted significantly to a linear model (Table 1), and the explained variance varied between 44 to 64 %. Latitude was related significantly only to the richness of SEED level 2 groups of genes (Table 1).

Finally, we also analyzed the taxonomic and functional diversity among types of habitat (or substrate) from which the sample was originally obtained (mat, water, or a miex of both). For both taxa and functions, the diversity was higher in mat, followed by water and both habitats pooled (Table 2). Using ANOVA, we found that the effect of substrate (mat, water, or mixed) on the logarithm of the number of taxa, genes, or groups of genes was significant in both taxa and functions (Table 2). Post-hoc comparisons showed significant differences among mat, water, and both types of substrates (Table S2).

**Table 2.** ANOVA results (η-squared and p-value) for comparisons of number of taxa and SEED 1, 2, and 3 categories among mat, water, and both (pooled).

| Relationship | η-squared | p-value |
| --- | --- | --- |
| Taxa-substrate | 0.33 | 0.04 |
| SEED 1-substrate | 0.61 | 0.0005 |
| SEED 2-substrate | 0.72 | 3.67e-5 |
| SEED 3-substrate | 0.67 | 0.0001 |

Mean and standard deviation (sd) values of number of taxa for mat, water and both were $102 \pm 40.9$ , $72.4 \pm 34.5$ , and $25 \pm 1.41$ . Mean and sd values of number of SEED 1 categories for mat, water and both were $39.1 \pm 1.05$ , $37.6 \pm 1.77$ , and $33 \pm 1.41$ . Mean and sd values of number of SEED 2 categories for mat, water and both were $560 \pm 31.1$ , $488 \pm 23.2$ , and $261 \pm 88.9$ . Mean and sd values of number of SEED 3 categories for mat, water and both were $426 \pm 20.1$ , $355 \pm 102$ , and $118 \pm 29.7$ . In all cases richness was higher mat followed by water and both.
$\eta$ (eta)-squared corresponds to the quotient of sum of squares of the model and the sum of squares of the total, i.e. the percent of explained variance

## Discussion

The importance of developing models in science lies in their objective capacity and predictive usefulness. The ideal is to generate models derived from primary principles, simple formulation (and few parameters), and that explain different variables from a single explanatory variable. The models that have these properties and wide empirical support have been called efficient theories, within which the metabolic theory of ecology is included (Marquet et al. 2014). In microbial ecology, in the last decade, macroecological approaches have been used, evaluating models originally thought for macroorganisms and developing alternatives with valid assumptions for microorganisms (Barberán et al. 2014). For example, in the MTE the assumption that in the temperature range 0-40°C the response is exponential (or linear on a logarithmic scale; Gillooly et al. 2001), an assumption that is not very well supported in microorganisms which have a curved behavior within this range or as the case of thermophiles and hyperthermophiles have a positive exponential growth outside this range (e.g. Sharp et al. 2014). Therefore, we derived a general model (Eq. 1.2, Chapter 1) applicable to plants, animals, and microorganisms and within the entire temperature range of life, but which was derived from primary principles and is simple (i.e., it only has three parameters and on a logarithmic scale can be adjusted using multiple linear regression). We used this general model to develop a theoretical framework for the structure of microbial metagenomes (Arroyo 2020, Chapter 2), which includes some of the predictions tested here.

We evaluated this microbial metagenomic framework in microbial communities from hot springs because these ecosystems are a good model to study temperature responses due to the wide temperature ranges at spatial scales of a few centimeters, e.g. 30-60°C; Mackenzie 2014, or meters (30-90°C, Sharp et al. 2014, Wang et al. 2014). Our results showed that, for both taxa, their metabolisms and genes, in the studied temperature range, there was a decline in diversity with temperature, with the tipping point being found before or slightly above 30°C. The value of the parameters varied from genes to metabolisms and species, as is expected in our metabolic ecological metagenomics framework (see Arroyo 2020, Chapter 2), the parameter values depend on the total number of reads sequenced, which increases when recruiting from genes to species.

Our approach differs from traditional studies in metagenomic and particularly in hot springs, which have often used comparative approaches to show that temperature is a variable that explains variation in both taxa and functions (in the mesophile-thermophile-hyperthermophile range; ∼30-90 ºC; Sharp et al. 2014). However, a few studies have tested predictions of ecological theories, such as Glazier et al. (2012), who showed that both the number of species and food chain length do not respond well to Arrhenius. Moreover, recent models have been developed in order to explain richness in these ecosystems (Klales et al. 2012).

The mechanisms behind these relationships depend on the spatio-temporal scale. At short spatiotemporal scales, they are mostly influenced by the number of individuals (MacArthur and Wilson 1967, Hubbell 2001) and genes (Vellend 2003), and their maintenance depends on different mechanisms (Chesson 2000, Chesson 2018), such as resource partitioning. Briefly, many populations with similar metabolisms can coexist in the same community if populations use different resources in space and/or time. Consequently, more productive patches can sustain more individuals, reducing the probability of extinction and concomitantly leading to more species, metabolisms, and genes (see Allen et al. 2007). At a large spatiotemporal scale, the effect of temperature on diversification rates (Allen et al. 2007) would also contribute to affecting the richness of taxa, and subsequently genes and metabolisms (Arroyo et al. 2019, Chapter 2). Our results show that the general principles that explain diversity in other ecosystems also apply to extreme temperature environments, and subsequently, the same models or theory can be used universally, irrespective of taxa. The fact that the pattern of temperature dependence for genes that we found in global hot springs is similar to that found in the global ocean (Sunagawa et al. 2015) and opposite to that found in soil may be due to the lower temperature range in soil ecosystems (Ding et al. 2015).

Temperature is not the only variable determining diversity in these ecosystems as we also showed that type of substrate, pH, and latitude can explain significant amounts of the variance, as also showed in previous studies (Miller et al. 2009, Ionescu et al. 2010, Cuecas et al. 2011, Tobler et al. 2011, Everroad et al. 2012, Cole et al. 2013, Wang et al. 2013, Inskeep et al 2013, Headd et al. 2014, Herbold et y. 2014, Valverde et al. 2014, Sharp et al. 2014, Wang et al. 2014; Menzel et al 2015; Power et al 2018). The significant effect of pH emphasizes the need to extend the metabolic theory of ecology to include effects of pH, as this chemical variable also influences enzymatic rate (Michaelis 1922, Alberty and Massey 1954; Alberty, 2006) and subsequently metabolism (Rektorschek et al. 1998), and other properties of ecological systems above the individual-level, such as microbial species richness (Fierer and Jackson 2006). The effects of pH are especially important in organisms that live embedded in substrates, such as those inhabiting soil and aquatic environments, including the ocean and freshwater habitats. Developing a mechanistic but simple model for the effects of pH is of applied importance to predict ecological responses to freshwater acidification, a task that often has been done using empirical approaches (e.g., Ou et al. 2015, Beaune et al. 2018). Further analysis distinguishing Bacteria and Archaea, including functions defined by other databases (e.g. InterPro2GO; Hunter et al. 2014, eggNOG; Powell et al. 2012) and with an increased number of subsamples and/or samples to achieve a higher statistical power, need to be carried out in other hotsprings in order to test the generality of the patterns herein reported.

In summary, our analyses showed that genes, metabolisms, and taxa can be well predicted by temperature, supporting the hypothesis that, given that temperature affects assemblage abundance, ultimately it should also affect the number of genes and metabolisms in a community. Moreover, our analysis also showed that pH explains a large amount of variance, indicating the need to extend ecological theory to include variables other than temperature.

## Methods

### Data and statistical analysis

Data of hot springs metagenomes consisted of metagenomes obtained from previous studies and metagenomes sampled and sequenced by us. We obtained and sequenced eight hot springs mat samples as described in (Guajardo-Leiva et al. 2018) for which the temperature and geographic coordinates were recorded. Additional metagenomic data were obtained from public databases from 18 other hot springs metagenomes spanning latitudinal (from Iceland to Antarctica) and longitudinal (from Yellowstone National Park in North America to Malaysia) geographic gradients, including six continents. Each of these samples was obtained from public databases and was obtained from either mat or water or a mix (of both mat and water; e.g., Menzel et al. 2015) and has a record of temperature and geographic coordinates. Our integrated database from our sampling and compiled metagenomic data encompasses a wide range of temperature (from 32 to 98°C), pH (from 2.5 to 9.25), and geographic distance (1 cm-19000 km). Metagenomic data includes different technologies of next-generation sequencing (NGS), including Illumina HiSeq, Ion Torrent, and Illumina MiSeq (Sanger and 454 were not included because of low depth of data; see Table S1). (However, to make these different technologies comparable, we apply a set of procedures, see below).

Raw metagenomic data were preprocessed with cutadapt version 1.8.1 (Martin 2011) to remove 5’ adapter and 3’ low-quality sequences. The metagenomic data that we have used were derived from three different technologies, which differ in read length and sequencing depth. It has been shown that different sequencing technologies can be comparable (Luo et al. 2012), however, in order to avoid errors, we made some corrections. To correct for differences in read length, we have established a minimum length of 30 bp, which is a plausible length for annotation (Carr and Borenstein 2014, Sims et al. 2014). All sequences with quality <30 were removed. To avoid biases in the estimation of diversity because of variable depths, after preprocessing, sequences were rarefied. To do this, it was randomly subsampled five million reads (the minimum depth among the 23 samples) in each sample using fastq-tools (https://homes.cs.washington.edu/~dcjones/fastq-tools/). Direct alignment of the preprocessed and subsampled nucleotide sequences against protein non-redundant NCBI database (NCBI-nr) was performed using DIAMOND (Buchfink et al. 2015) with an e-value of 1e-10. To overcome differences in sequencing deep not effect because we have randomly subsampled the same amount of reads among samples. The taxonomic and functional assignments based on the alignments were interpreted using MEGAN6 (Huson et al. 2016). Here we based on functions assigned by the SEED database (Overbeck et al. 2005, Overbeck et al. 2014). Briefly, this database of genome annotation employs the subsystem approach. A subsystem is a set of functional role that constitutes a biological process or structure. Then a subsystem includes metabolic pathways (glycolysis) and other protein interactions such as ribosomes. The classification of genes within subsystem is hierarchically nested, where in the lowest level (SEED level 1) includes the more general functions with 44 categories (e.g. membrane transport, motility and chemotaxis, stress response, among others), the intermediate level of resolution includes nested functions (SEED level 2, 1136 categories; e.g. glutamate dehydrogenases, glycine biosynthesis, etc.), and the highest level correspond to genes (SEED level 1, 5738 categories; e.g. arginase, cysteine synthase A, etc.).

Relationships between the number of taxa, genes, and groups of genes and temperature were performed using (simple or multiple) linear regression, according to the Eyring-Polanyi eq. of two parameters (eq. 1.1) or eq. 1.2. Regressions were performed using the lm function as implemented in the R programming environment. Empirical relationships between the logarithm of the number of taxa, genes, and metabolisms and pH, and latitude were assessed using simple linear regression. Also, using the lm function. To test the effect of the type of substrate (mat, water or both) we performed ANOVA using the aov and lm functions as implemented in R.

## Acknowledgments

JIA acknowledges support from a CONICYT National Doctoral Scholarship N° 21130515

## Notes

### Competing Interest Statement

The authors have declared no competing interest.

