## Supplementary Material for "Global patterns of taxonomic and functional diversity in hot springs microbiomes"

José Ignacio Arroyo[1]* Beatriz Díez[2] and Pablo A. Marquet[1, 3, 4, 5, 6, *]
1. Departamento de Ecología, Facultad de Ciencias Biológicas, Pontificia Universidad Católica de Chile
2. Departamento de Genética Molecular y Microbiología, Facultad de Ciencias Biológicas, Pontificia Universidad Católica de Chile, CP 8331150, Santiago, Chile.
3. Instituto de Ecología y Biodiversidad (IEB), Las Palmeras 3425, Santiago, Chile.
4. Laboratorio Internacional de Cambio Global (LINCGlobal) and Centro de Cambio Global UC, Facultad de Ciencias Biológicas, Pontificia Universidad Católica de Chile, CP 8331150, Santiago, Chile.
5. The Santa Fe Institute, 1399 Hyde Park Road, Santa Fe NM 87501, USA.
6. Instituto de Sistemas Complejos de Valparaíso (ISCV), Subida Artillería 470, Valparaíso, Chile.

*Corresponding authors:
José Ignacio Arroyo,
Pablo Marquet,

### Supplementary tables

Table S1. List of sequenced and compiled metagenomic data. Numbers of samples were defined from north to south and west to east.

| Sample ID | | T (°C) | pH | Geographic coordinates | Accession number |
|---|---|---|---|---|---|
| Our laboratory | 82TAT | 82 | 6 | -22.32, -68.009 | G1A* |
| | 60TAT | 60 | 6.8 | -22.32, -68.008 | G2D* |
| | 44K | 44 | 6.1 | -62.98, -60.58 | K1A* |
| | 32K | 32 | 6.2 | -62.98, -60.58 | K2A* |
| | 42P | 42 | 9.25 | -42.25, -72.36 | C3* |
| | 48P | 48 | 7.1 | -42.45, -72.46 | P4* |
| | 58P | 58 | 6.9 | -42.45, -72.46 | P3* |
| | 66P | 66 | 6.8 | -42.45, -72.46 | P1* |
| Menzel | 65C | 65 | 7 | 26.25, 99.99 | Ch2-EY65S |
| Menzel | 90IC | 90 | 3.75 | 63.90, -22.05 | 4530144.3§ |
| Menzel | 76IT | 76 | 3 | 40.82, 14.13 | Is3-13 |
| Lin et al. 2015 | 69TAI | 69 | 2.5 | 25.19, 121.60 | 4583585.3§ |
| López-López et al. 2015 | 76S | 76 | 8.2 | 41.86, −8.10 | It6 |
| Saxena et al. 2017 | 55IN | 55 | 7.8 | 22.65, 78.36 | 4529716.3§ |
| | 43.5IN | 43.5 | 7.5 | 22.65, 78.36 | SRR1297204 |
| | 52.1IN | 52.1 | 7.8 | 23.41, 83.39 | SRR2239652 |
| | 61.5IN | 61.5 | 7.6 | 23.41, 83.39 | SRR3961733 |
| | 69IN | 69 | 7 | 23.41, 83.39 | SRR3961734 |
| | 67IN | 67 | 7.8 | 23.41, 83.39 | SRR3961739 |
| https://www.ncbi.nlm.nih. | 60YNP | 60 | 8 | 44.555, -110.835 | SRR3961741 |
| gov | | | | | SRR3961742 |
| /sra/?term=SRR5248366 | | | | | SRR3961743 |
| | | | | | SRR5248366 |

Value which precedes the sample ID correspond to temperature

| Table S2. Post hoc Tukey comparisons among mat, water, and both. | | |
|---|---|---|
| Relationship | Comparison | P-value |
| Taxa-Substrate | Mat-Both | 0.04 |
| SEED 1-Substrate | Mat-Both<br>Water-Both | 0.0001<br>0.002 |
| SEED 2-Substrate | Mat-Both<br>Water-Both<br>Water-Mat | 0.00003<br>0.0007<br>0.06 |
| SEED 3-Substrate | Mat-Both<br>Water-Both | 0.00009<br>0.001 |
| η (eta)-squared corresponds to the quotient of sum of squares of the model and the sum of squares of the total, i.e. the percent of explained variance | | |
